# LieMap: A Fast and Transparent Gradient-Based Method for Cryo-EM Structure Fitting

**DOI:** 10.1101/2025.09.16.676663

**Authors:** Yue Hu, Zanxia Cao, Yingchao Liu

## Abstract

Fitting high-resolution atomic models into lower-resolution cryo-electron microscopy (cryo-EM) maps is a critical yet challenging step in structural biology. We present LieMap, a straightforward, fast, and transparent gradient-based fitting protocol designed for practical applications. Our method parameterizes rigid-body motion on the Lie algebra `se`(3) to provide a minimal and singularity-free representation. The optimization objective is to directly maximize a real-space cross-correlation score, calculated as the mean density value sampled from the map at the atomic coordinates. This score is computed differentiably using trilinear interpolation and optimized with the Adam optimizer in a multi-stage fashion. We demonstrate the effectiveness of LieMap by fitting the apo-state structure of *E. coli* GroEL (PDB: 1aon) into its ATP-bound state cryo-EM map (EMD-1046). The protocol successfully navigates the large conformational change, achieving a final RMSD of 3.23 Å. LieMap’s simplicity, computational efficiency, and transparent optimization process make it a valuable and practical tool for structural biologists.

## 1 Introduction

The integration of high-resolution atomic models with cryo-EM density maps is a cornerstone of modern structural analysis [1]. This process, known as fitting, aims to find the optimal position and orientation of a known atomic structure within the 3D map. The primary challenges in this rigid-body docking problem are the vastness of the six-dimensional search space and the often complex, multi-modal nature of the scoring landscape.

Historically, this challenge has been addressed by a variety of methods. Exhaustive 6D cross-correlation searches, while robust, are computationally prohibitive for routine tasks. Faster methods based on the Fast Fourier Transform (FFT) excel at translational searches but require a separate, often discretized, rotational search. Other methods may rely on stochastic optimization, like Monte Carlo, which can be slow to converge. A common limitation of these approaches is their ”black-box” nature, where the user has little insight into the optimization trajectory until the process is complete.

The rise of deep learning frameworks has popularized the use of gradient-based optimization. To leverage these powerful techniques, every component of the fitting pipeline must be differentiable. Here, we present LieMap, a method that embraces this paradigm for its simplicity, speed, and transparency. It combines three key ideas:

1. A singularity-free representation of rigid motion using the Lie algebra `se`(3).
2. A direct, differentiable cross-correlation score based on trilinear interpolation.
3. A multi-stage optimization protocol using the Adam optimizer, which is fast, robust, and provides transparent monitoring of the fitting process.

This paper details the mathematical and algorithmic implementation of LieMap and demonstrates its practical utility on a challenging, real-world test case involving the GroEL/GroES chaperonin system [4].

## 2 Methods

The core of the LieMap algorithm is to iteratively update the pose of a mobile atomic structure, represented as a point cloud 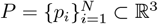, to maximize its overlap with a target density map *ρ*: ℝ^3^ → ℝ.

### 2.1 Data Preparation

Before fitting, both the atomic model and the density map are pre-processed. The atomic coordinates are parsed from standard PDB or mmCIF files. The cryo-EM map is loaded from an MRC file, and a binary mask is generated by applying a sigma-level threshold (typically 3.0*σ*). This crucial step focuses the optimization on the highest-confidence regions of the density map, effectively reducing the influence of background noise. For the initial coarse-grained search stage, this masked map is further processed with a Gaussian blur (sigma=4.0). This blurring smooths the scoring landscape, removing small, noisy local maxima and creating a broader, more convex-like basin for the global minimum, which helps prevent the optimizer from getting trapped prematurely.

### 2.2 Rigid Motion on the Lie Algebra

A rigid-body transformation in 3D space is an element of the Special Euclidean group, SE(3). Instead of parameterizing this transformation with Euler angles, which suffer from singularities (gimbal lock), we use the corresponding Lie algebra `se`(3), a concept well-established in fields like robotics [3]. An element of `𝔰𝔢`(3) is a 6-dimensional vector *ξ* = (*ω, v*) ∈ ℝ^6^, often called a ”twist”. The first three components, *ω* = (*ω*_*x*_, *ω*_*y*_, *ω*_*z*_), define the axis and angle of rotation, while the last three, *v* = (*v*_*x*_, *v*_*y*_, *v*_*z*_), define the translation vector.

This 6-parameter vector provides a minimal, unconstrained, and continuous representation of the transformation. At each optimization step, the optimizer can freely update these six values. These values are then mapped back to a valid transformation matrix. The rotational part is recovered by first forming the 3 × 3 skew-symmetric matrix 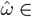 `𝔰𝔬`(3):

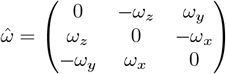

and then applying the matrix exponential to yield the rotation matrix *R* ∈ SO(3):

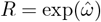

This exponential map ensures that any update to *ω* results in a valid rotation matrix. The final transformation of a point *p*_*i*_ to its new position 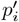 is given by:

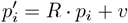

### 2.3 Differentiable Scoring and Optimization

To guide the optimization, we use a direct, real-space density score. For each transformed atomic coordinate 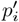, we sample the density value from the map *ρ*. Since 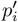 has continuous coordinates, its value on the discrete voxel grid of the map is calculated using trilinear interpolation. The final fitting score, *S*, is the mean density value sampled from the map at the coordinates of all *N* atoms. This serves as a direct cross-correlation score, where higher density values correspond to a better fit:

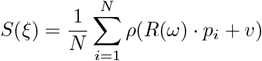

Our goal is to find the parameters *ξ*^*^ that maximize this score. This is achieved by minimizing the negative score, *L*(*ξ*) = − *S*(*ξ*), using the Adam optimizer [2]. The gradient of the loss with respect to the six parameters of *ξ* is computed efficiently via automatic differentiation, a core feature of modern deep learning frameworks like PyTorch.

### 2.4 The Multi-Stage Optimization Strategy

To balance speed and accuracy, we employ a coarse-to-fine, multi-stage protocol:

1. **Pre-alignment:** A trivial initial translation is applied to align the center of mass of the atomic structure with the center of mass of the density map mask. This places the structure in the correct vicinity to start the search.
2. **Stage 1: Coarse-grained Search:** A rapid, coarse search is performed for 500 steps with a relatively high learning rate (lr=0.01). This stage uses only the C-alpha atoms and the blurred density map. Using a subset of atoms reduces the computational cost, while the blurred map, as mentioned, helps locate the global minimum’s basin.
3. **Stage 2: Fine-grained Refinement:** A much longer refinement is performed for 5000 steps with a smaller learning rate (lr=0.005). This stage uses all atoms of the structure and the original, non-blurred density map. The increased number of points and the high-resolution map allow the optimizer to make fine adjustments and settle into a precise final pose.

## 3 Results and Discussion

We tested LieMap on the challenging case of fitting the apo-state GroEL structure (PDB: 1aon) into the cryo-EM map of its ATP-bound state (EMD-1046) [4]. This involves a significant conformational change. The final fit was validated against the refined gold-standard structure (PDB: 1GRU).

### 3.1 Performance, Transparency, and Practicality

A key advantage of LieMap is its exceptional speed and transparency. The entire two-stage optimization process completed in approximately **18.3 seconds** on an Apple M4 mini platform. This efficiency is critical for high-throughput structural analysis and rapid hypothesis testing.

Furthermore, the method provides a ”glass-box” view of the optimization. The score and RMSD are monitored at every 10th step, offering immediate, real-time feedback. This transparency is a significant practical advantage over ”black-box” methods. It allows a user to instantly assess the fitting trajectory and diagnose potential issues. For example, a stagnating or decreasing score is a clear indicator that the learning rate may be too high or that the optimizer is trapped, allowing the user to intervene, adjust hyperparameters, and restart without wasting significant time.

### 3.2 Fitting Accuracy

The optimization protocol was highly effective. The key metrics of the fitting process are summarized in Table 1. The coarse-grained stage rapidly found the correct basin, reducing the RMSD from an initial 14.93 Å to 9.82 Å. The subsequent fine-grained refinement steadily improved the fit to a final RMSD of **3.23 Å**. The visual result of this process, generated with PyMOL [5], is depicted in Figure 1. The excellent visual overlap between our fitted model and the gold-standard structure confirms the high accuracy of the result.

**Table 1:** Summary of the LieMap fitting results for PDB 1aon into map EMD-1046.

| Metric | Value |
| --- | --- |
| Initial RMSD (Post-CoM) | 14.93 Å |
| RMSD after Stage 1 (Coarse) | 9.82 Å |
| Final RMSD (after Stage 2) | 3.23 Å |
| Total Execution Time | 18.3 seconds |

**Figure 1.**
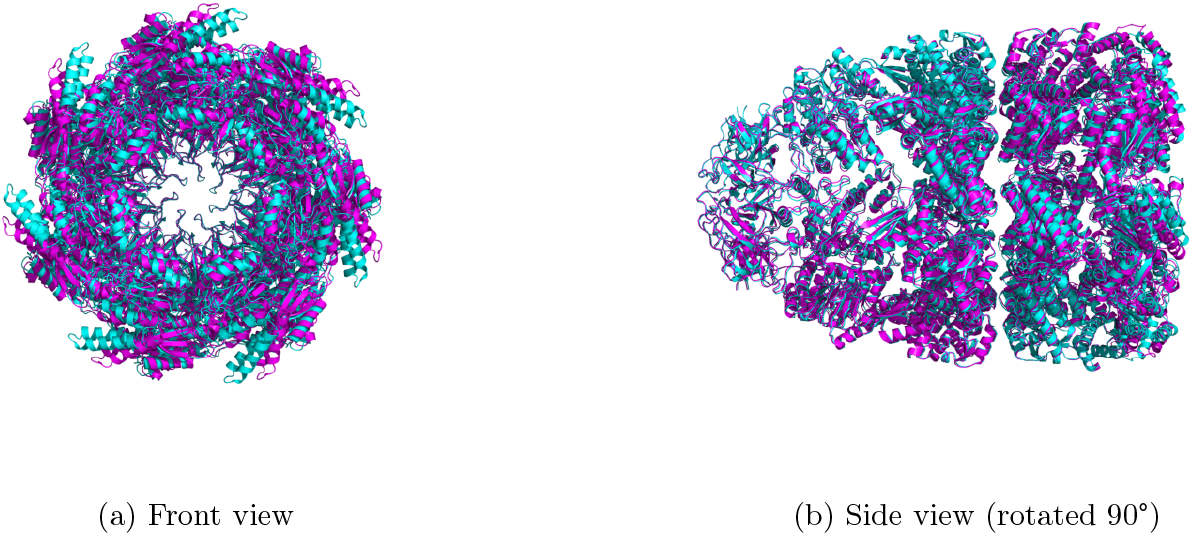
Superposition of the final fitted model (1aon, colored cyan) and the gold-standard structure (1GRU, colored magenta). The close overlap from multiple viewpoints demonstrates the high accuracy of the fit.

### 3.3 Limitations and Future Directions

Despite its success, LieMap has limitations inherent to its design as a local optimizer. Its performance is dependent on the initial pre-alignment placing the model reasonably close to the correct solution. It is not designed for a global search from an arbitrary starting position. Future work could address this by integrating a coarse global sampling step (e.g., a low-discrepancy rotational search) before initiating the gradient-based refinement.

Additionally, the current implementation performs a rigid-body fit. Many biological processes involve flexible conformational changes. A future direction would be to extend LieMap to support flexible fitting, for instance by defining multiple rigid bodies connected by flexible linkers or by incorporating deformations along normal modes.

## 4 Conclusion

LieMap demonstrates that a combination of well-established mathematical and computational tools can lead to a powerful and practical solution for cryo-EM fitting. By using a Lie algebra parameterization for transformations and a direct, differentiable density score, we unlock the ability to use modern gradient-based optimizers like Adam effectively. The resulting method is fast, transparent, and accurate, achieving a 3.23 Å RMSD on a difficult test case. Its simplicity and interpretability make it an excellent tool for the practicing structural biologist.

## Code Availability

The source code for LieMap is publicly available at https://github.com/YueHuLab/LieMap.

## Acknowledgments

The human authors designed the entire algorithm and are responsible for the results and the process. We thank Gemini for its assistance in code implementation and document preparation.

## References

[1] Frank, J. (2006). Three-dimensional electron microscopy of macromolecular assemblies: visualization of biological molecules in their native state. Oxford university press.

[2] Kingma, D. P., & Ba, J. (2014). Adam: A method for stochastic optimization. arXiv preprint 1412.6980.

[3] Murray, R. M., Li, Z., & Sastry, S. S. (1994). A mathematical introduction to robotic manipulation. CRC press.

[4] Ranson, N. A., Farr, G. W., Roseman, A. M., Gowen, B., Fenton, W. A., Horwich, A. L., & Saibil, H.R. (2001). ATP-bound states of GroEL captured by cryo-electron microscopy. Cell, 107(7), 869–879.

[5] The PyMOL Molecular Graphics System, Version 3.1, Schrödinger, LLC.

